# Position-sensing established during compaction dictates cell fate in the mammalian embryo

**DOI:** 10.1101/856161

**Authors:** Christophe Royer, Karolis Leonavicius, Annemarie Kip, Deborah Fortin, Kirtirupa Nandi, Anna Vincent, Celine Jones, Tim Child, Kevin Coward, Chris Graham, Shankar Srinivas

## Abstract

Precise patterning within the 3-dimensional context of tissues, organs and embryos implies that cells can sense their relative position. We still understand relatively little about how cells sense their relative position within small groups of cells. During preimplantation development, outside and inside cells rely on apicobasal polarity and the Hippo pathway to choose their fate. Despite recent findings suggesting that mechanosensing may be central to this process, the relationship between blastomere geometry (i.e. shape and position) and the Hippo pathway effector YAP remains unknown. To address this, we used a highly quantitative approach to collect and analyse information on the geometry and YAP localisation of individual blastomeres of mouse and human embryos. We find that the fraction of YAP in the nucleus responds to the proportion of exposed cell surface area rather than blastomere shape. To directly test the influence of blastomere position on YAP localisation we developed an approach using hydrogel based cylindrical cavities to alter blastomere arrangement in cultured embryos. Unbiased clustering analyses of blastomeres from such embryos reveal that the ability to respond to position emerges as early as the 8-cell stage during compaction and that altering the relative position of a blastomere within an embryo can alter its fate. Finally, we demonstrate that this response is lost upon inhibiting PKC signalling, linking it to the cell polarity. Our results pinpoint when during early embryogenesis cell specification decisions are initiated and show how blastomeres can sense their position in a polarity dependent manner to alter YAP localisation.

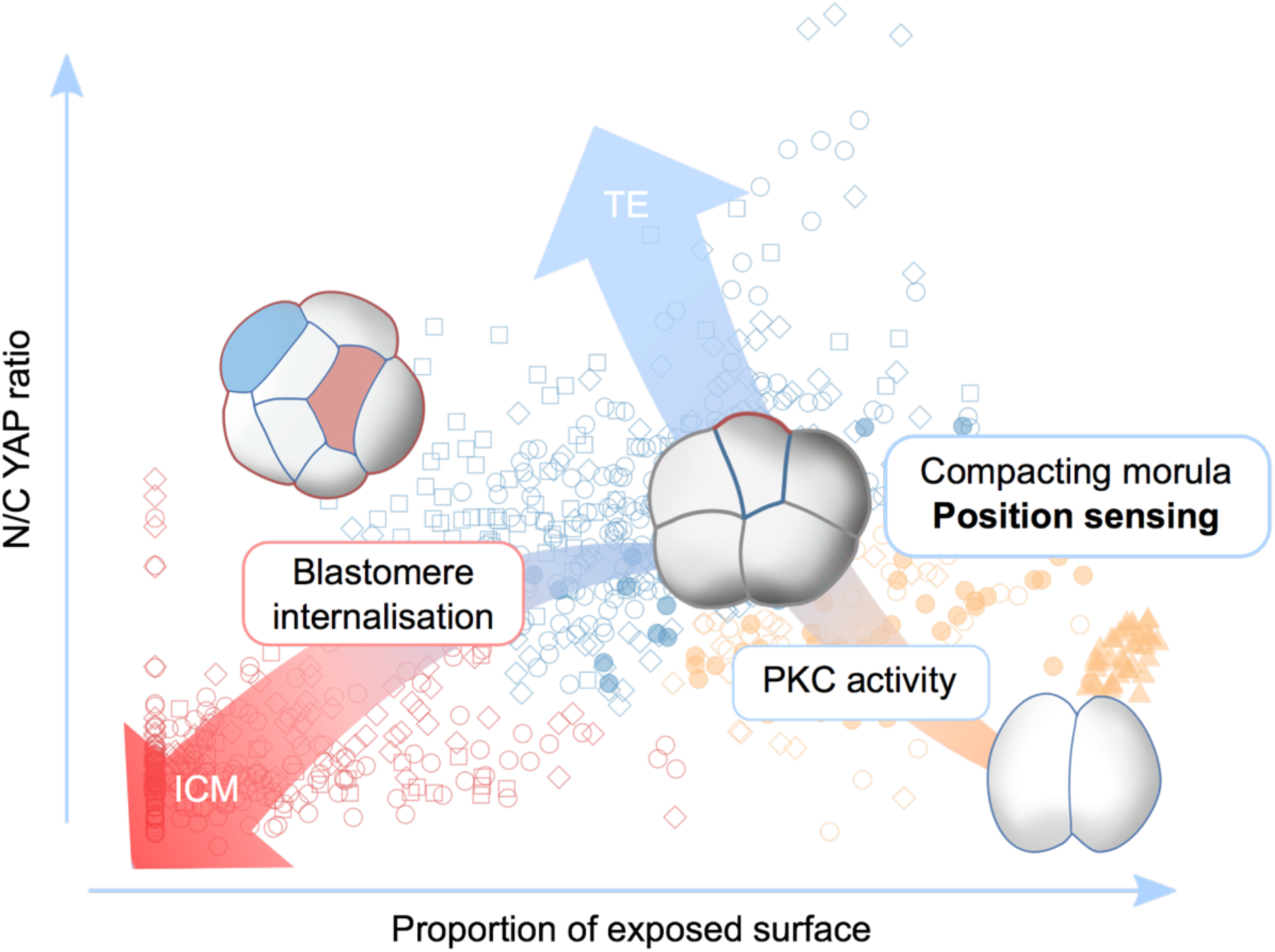

## INTRODUCTION

Through the course of embryonic development, complex tissues and organs are formed from a single cell. This requires each given cell within the developing embryo to constantly sense its position within its 3D environment in order to make the correct fate decisions. Morphogen gradients have long been suggested to play an important role in transmitting positional information across tissues during embryo patterning (Briscoe and Small, 2015; Tickle et al., 1975) though even here, mechanistic details remain unclear (Wolpert, 2016). When small groups of cells are concerned, morphogen gradients are conceptually harder to set up, leading cells to rely on other strategies. In the mouse pre-implantation embryo, around the 16-cell stage, blastomeres differentiate into trophectoderm (TE) or inner cell mass (ICM) cells depending on their outside or inside position respectively. The molecular mechanisms by which outside and inside cells determine their fate has been extensively studied, highlighting the importance of apicobasal polarity and Hippo signalling (Plusa et al., 2005; Sasaki, 2017; White et al., 2018).

In apolar inner cells, AMOT localises at cell-cell junctions where it associates with NF2 and is phosphorylated by the Hippo pathway kinases Lats1/2 (Cockburn et al., 2013; Hirate et al., 2013). In concert with Lats1/2, AMOT is then able to Induce the phosphorylation of YAP and its sequestration to the cytoplasm. In inner cells, YAP is consequently unable to bind to TEAD4 and cannot induce the transcription of TE genes such as *CDX2* (Nishioka et al., 2009, 2007). Instead, genes associated with pluripotency, such as *SOX2*, drive ICM fate in these cells (Wicklow et al., 2014). In outer cells, the establishment of apicobasal polarity generates a contact-free membrane domain where cortical F-actin sequesters AMOT away from cell-cell junctions, preventing its phosphorylation and interaction with Hippo pathway components (Hirate and Sasaki, 2014). Following up on a study on the role of Rho-ROCK signalling in TE polarisation and Hippo inactivation (Kono et al., 2014), it was shown that the interaction of AMOT with cortical F-actin is promoted by the activity of Rho which acts by preventing the phosphorylation of AMOT at S176 and the interaction between AMOT and NF2 (Shi et al., 2017). This results in the inability of the Hippo pathway to phosphorylate YAP, which can then translocate to the nucleus and interact with TEAD4 to drive the expression of TE-specific genes such as CDX2, consequently inducing TE fate. Together, this body of work, in addition to highlighting the importance of the Hippo pathway and apicobasal polarity in the first cell fate decision, also suggests multiple links between cytoskeletal organisation and Hippo pathway regulation.

It is now widely accepted that forces, related to cell shape changes or cell position within a group of cells, can modulate the localisation and activity of YAP via the actin cytoskeleton (Aragona et al., 2013; Dupont et al., 2011; Halder et al., 2012; Wada et al., 2011). In the preimplantation embryo, although it has been suggested that mechanosensing may occur (Maître et al., 2016) and despite the multiple links between the Hippo pathway and the actin cytoskeleton, it remains unclear whether the shape or position of individual blastomeres can directly regulate the subcellular localisation of YAP to modulate cell fate. To answer this question, we generated a collection of embryos from the 2- to the 64-cell stage, analysing in conjunction the subcellular localisation of YAP and individual blastomere geometry. We used a multivariate analysis including blastomere shape and position descriptors within the embryo to show that the proportion of exposed blastomere surface area, rather than shape has the strongest association with differences in YAP localization. Through unbiased clustering analysis and using non-invasive methods to modulate overall embryo geometry, we also find that the relationship between blastomere position and YAP subcellular localisation emerges as early as the 8-cell stage, during compaction. Finally, we show that the ability of a blastomere to sense its position via its proportion of exposed surface area ultimately determinates its fate.

## RESULTS

### Cells with high N/C YAP ratio occur prior to the first cell fate decision

In a 2D environment, cellular parameters such as shape and position, that we refer to as cell-geometry, have been shown to affect the localisation (and therefore activity) of YAP through its mechanosensing properties (Aragona et al., 2013; Dupont et al., 2011; Wada et al., 2011). In the pre-implantation embryo, despite the suggestion that mechanosensing may be involved in the regulation of YAP activity (Maître et al., 2016), it remains unclear whether geometrical properties of blastomeres directly influence the relative distribution of YAP to the cytoplasm and nucleus. To analyse the relationship between the localisation of YAP and the geometry of individual blastomeres during mammalian preimplantation cell fate allocation, we used a highly quantitative approach combining imaging, manual segmentation and image analysis. We collected and stained mouse embryos for YAP, F-actin and E-cadherin at the 2- (n=20), 8- (n=10), 16- (n=17), 32- (n=12) and 64-cell (n=2) stages (Figure 1A). We next created 3D cellular-resolution volume representations of these embryos by manually segmenting cell membrane and nucleus of individual blastomeres (Figure 1B and movie S1). This allowed us to accurately quantitate the cellular localisation of YAP in the nuclear and cytosolic compartments, by determining the ratio of nuclear to cytoplasmic YAP (N/C YAP ratio), and various geometric properties of blastomeres including, for example, shape factor and sphericity. We were also able to quantify the area of the blastomere limiting membrane that is exposed to the outside, in contact with other blastomeres and can be considered junctional (Leonavicius et al., 2017). This allowed us to determine the relative position of each blastomere within the embryo using the proportion of exposed surface area as a measure of whether a blastomere is embedded within the embryo or is on the surface.

**Figure 1:**
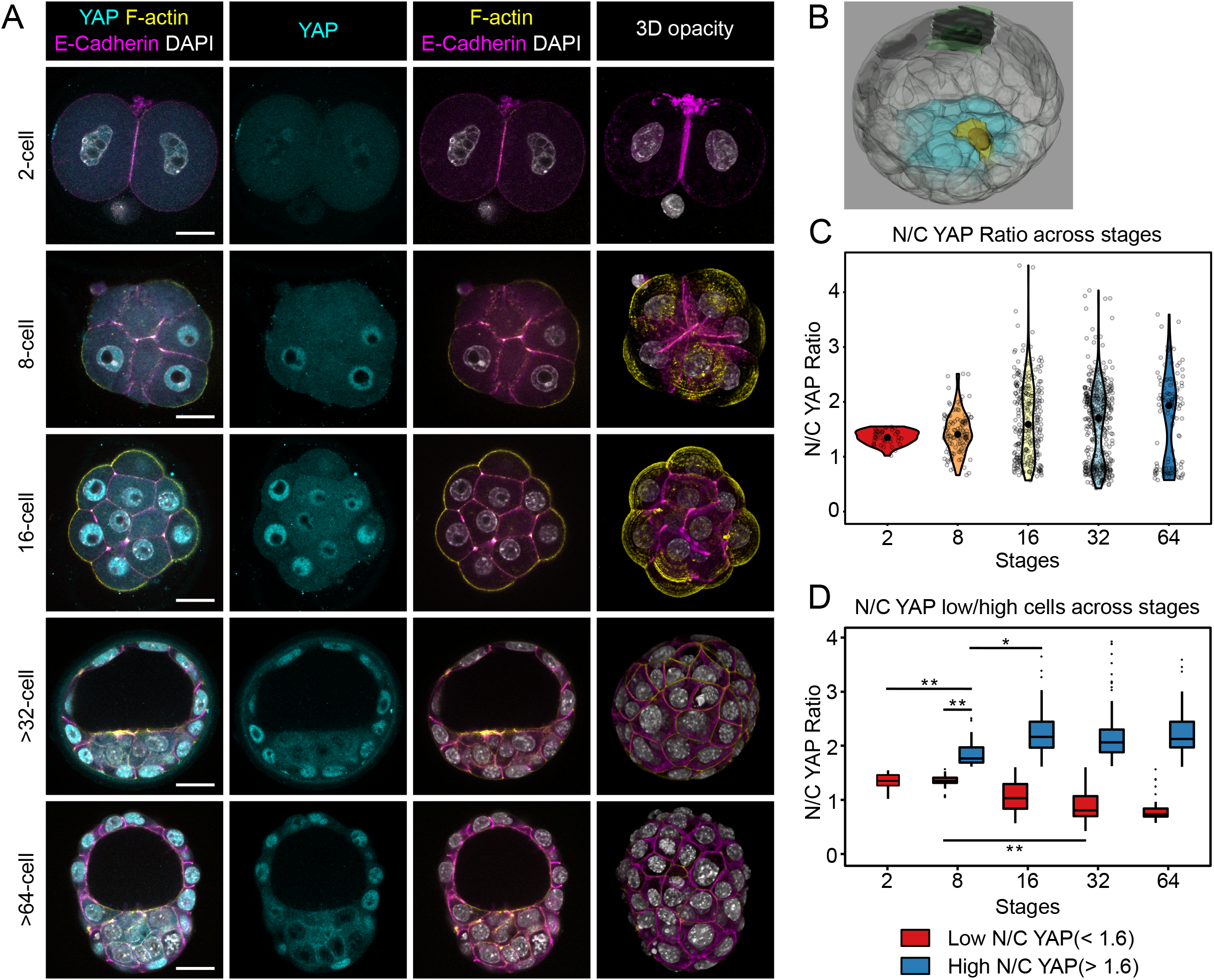
Analysis of N/C YAP ratio across preimplantation development using manual segmentation. A. Immunostaining of embryos at the 2- (n=20), 8- (n=10), 16- (n=17), 32- (n=12) and 64-cell stage (n=2) using antibodies against YAP and E-cadherin. F-actin and nuclei were visualised using Phalloidin and DAPI respectively. B. Example of a manually segmented 32-cell blastocyst showing blastomeres (green and yellow cells) exhibiting different shapes. Part of the cells making the trophectoderm are not displayed to be able to see inside the blastocyst cavity. The ICM is highlighted in cyan. C. N/C YAP ratio across developmental stages. Using membrane and nuclei volumes obtained from individual blastomeres, we were able to accurately extract information about the localisation of YAP by measuring N/C YAP ratio at the indicated stages of development. The mean N/C YAP ratio for each developmental stage is represented as a black dot. D. Low and High N/C YAP ratio blastomeres across developmental stages. Blastomeres were classified as exhibiting either high (>1.6) or low (<1.6) N/C YAP ratio based on K-means algorithm to separate them into two populations in an unbiased manner. Scale bar: 20 μm. NS: not significant. * p<0.05, ** p<0.01 (Kruskal Wallis test followed by Dunn’s test).

When we considered all blastomeres at each stage we found that, as expected, two populations of blastomeres with distinct N/C YAP ratios seemed to progressively appear from the 16-cell stage onwards (Figure 1C). In order to confirm this observation, we unbiasedly separated cells with high and low N/C YAP ratio using K-means clustering (Figure S1), that allowed us to define a N/C ratio of 1.6 as the threshold separating ‘low’ and ‘high’ nuclear YAP cells. Using this classification, we found that, as published before when comparing inside and outside cells, those two populations of cells become strongly separate from the 16-cell stage onwards (Figure 1D) (Hirate et al., 2015; Nishioka et al., 2009), thereby validating the representativeness of our data-set. However, our detailed quantification further revealed that these two populations could already be detected to be diverging at the 8-cell stage, suggesting that cells with relatively high N/C YAP ratio start arising even before outside and inside cells exist (Figure 1D).

### Blastomere position, as opposed to shape is strongly associated with N/C YAP ratio

To determine whether N/C YAP ratio corelated with any physical characteristics of blastomeres, we interrogated our quantitative data-set for the correlation between N/C YAP ratio and parameters describing blastomere shape (sphericity, oblateness, prolateness and volume) and position (proportion of exposed surface area and its converse, proportion of contact area). The proportion of exposed surface area and sphericity showed the strongest correlation with N/C YAP ratio (Figure 2A), so going forward we focused on these two parameters.

**Figure 2:**
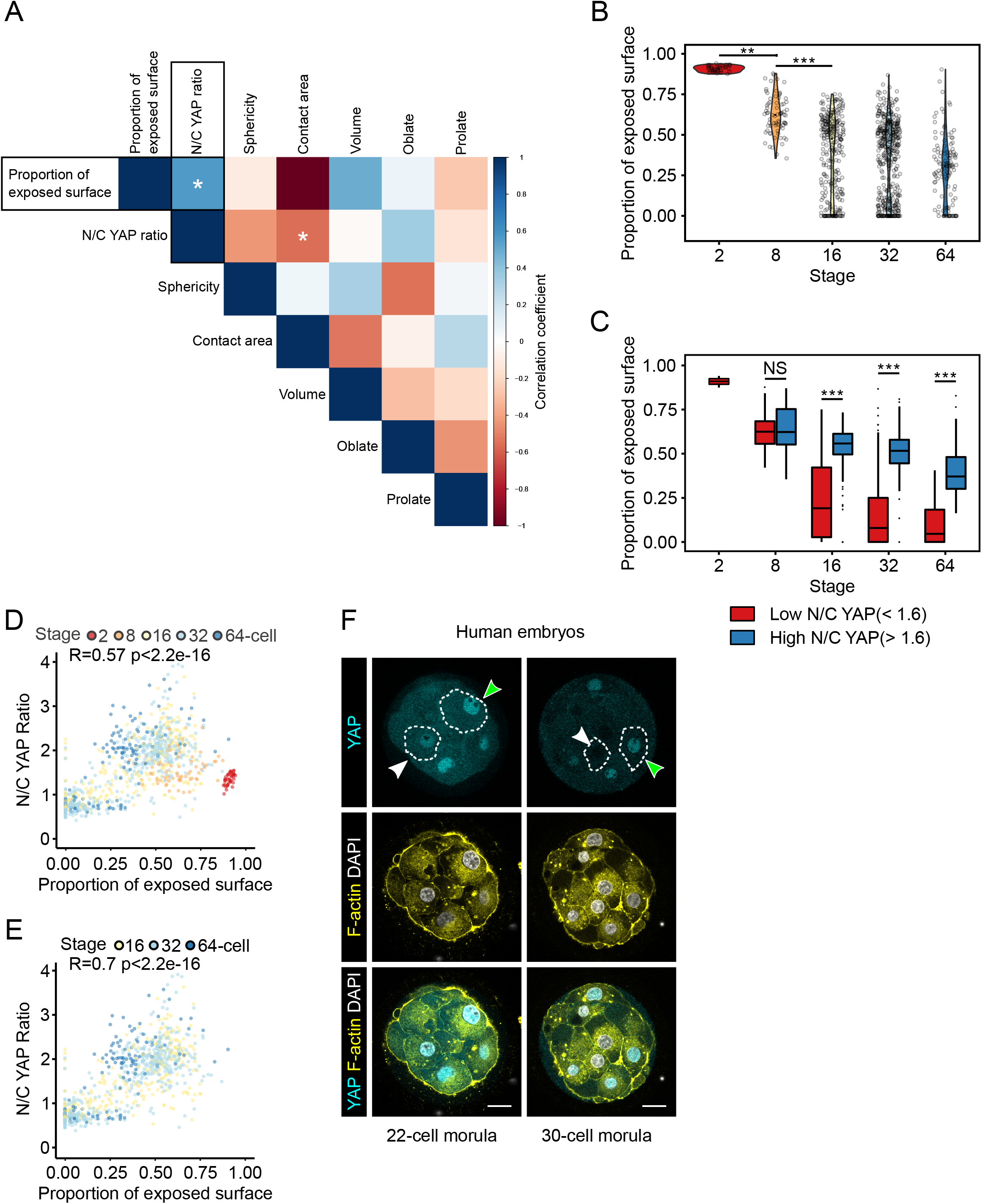
The proportion of exposed surface is associated with the proportion of YAP in the nucleus. A. Correlation matrix between N/C YAP ratio and geometry variables of individual blastomeres across preimplantation development. Note how the proportion of exposed surface and its converse, the proportion of contact surface correlate the highest with N/C YAP ratio. B. Proportion of exposed blastomere surface area across developmental stages. The mean proportion of exposed surface area for each developmental stage is represented as a black dot. C. Proportion of exposed cell surface area of blastomeres with either low or high N/C YAP ratio (as classified by K-means clustering) across developmental stages. NS: not significant. ** p<0.01, *** p<0.001 (Kruskal Wallis test followed by Dunn’s test). D. Correlation analysis between the proportion of exposed cell surface area (an indicator of position) and N/C YAP ratio throughout preimplantation development (Spearman, R=0.57 p<2.2e−16). E. Correlation analysis between the proportion of exposed cell surface area and N/C YAP ratio from the 16-cell stage onwards during preimplantation development (Spearman, R=0.7 p<2.2e−16). F. Representative optical sections of human morulae containing the indicated number of cells and immunostained for YAP. White arrowheads point at cells with either low or no exposed cell surface area and low nuclear YAP, whereas green arrowheads point at cells with high exposed cell surface area and high nuclear YAP. F-actin and nuclei were visualised using Phalloidin and DAPI respectively. Scale bar: 20 μm.

Sphericity decreased gradually from the 2- to the 64-cell stage, most likely because of the increasingly varied blastomere shapes arising as the embryos developed (Figure S2A). Though we could detect a statistically significant difference in sphericity of high and low N/C YAP ratio at the 16-cell stage, it was only at the 32-cell stage that the difference in median sphericity became really pronounced (Figure S2B). Furthermore, N/C YAP ratio and sphericity showed a relatively weak negative correlation (R= −0.39; p<2.2e−16) when analysing all blastomeres from the 2- to the 64- cell stage (Figure S2C). When only blastomeres of the 32- and 64-cell stage were considered (i.e. when outside cells become more stretched and elongated in shape), the association between N/C YAP ratio and sphericity was stronger, showing a moderate negative correlation (R= −0.54; p<2.2e−16) (Figure S2D). Together these data show that blastomere shape, based on sphericity, is a relatively poor predictor of YAP localisation, suggesting that it is unlikely that cell shape would directly modulate YAP localisation.

Unsurprisingly, we found that the proportion of exposed cell surface area continually decreased from the 2- to 64- cell stage, in line with the fact that as the ICM forms, more cells end up inside. We also saw a dramatic decrease in the proportion of exposed cell surface area from the 2- to 8-cell stage, highlighting the process of compaction (Figure 2B). Interestingly, cells with high and low N/C YAP ratios exhibited dramatic differences in the proportion of exposed cell surface area as early as the 16-cell stage (Figure 2C). We examined the relationship between the proportion of exposed cell surface area and N/C YAP ratio considering blastomeres across all stages and found that they are only moderately correlated (R=0.57; p<2.2e−16) (Figure 2D). However, this association became strong when considering blastomeres from the 16-cell stage onwards (R=0.7; p<2.2e−16) (Figure 2E).

To determine if a similar relationship existed in human embryos, we examined the localisation of YAP in human compacted morulae. We found that blastomeres with a high proportion of exposed surface area exhibited high levels of YAP in their nuclei, whereas cells on the outside that were more embedded within the embryo had reduced nuclear YAP and completely inside cells had close to no nuclear YAP. This suggests that cell fate allocation in the human follows the same principles as during mouse preimplantation development (Figure 2F). Together, our findings suggest that the amount of YAP in the nucleus is proportional to the extent to which cells are outside or inside, based on the proportion of exposed surface. This suggests that the molecular mechanism acting to regulate the localisation of YAP is able to very accurately sense the relative amount of exposed cell surface area.

### The relationship between the proportion of exposed surface and N/C YAP ratio is established during compaction

Though the relationship between the proportion of exposed cell surface area and N/C YAP ratio became strong only from the 16-cell stage onwards, it was not uncommon to see blastomeres at the 8-cell stage with a lower amount of YAP in the nucleus that also seemed to be more embedded within the embryo (Figure 3A). Furthermore, the range of N/C YAP ratio values at the 8-cell stage was drastically wider in comparison to the 2-cell stage, with cells exhibiting high and low N/C YAP ratio (Figure 1C and 1D). This suggests that blastomeres may already be able to sense their position at the 8-cell stage.

**Figure 3:**
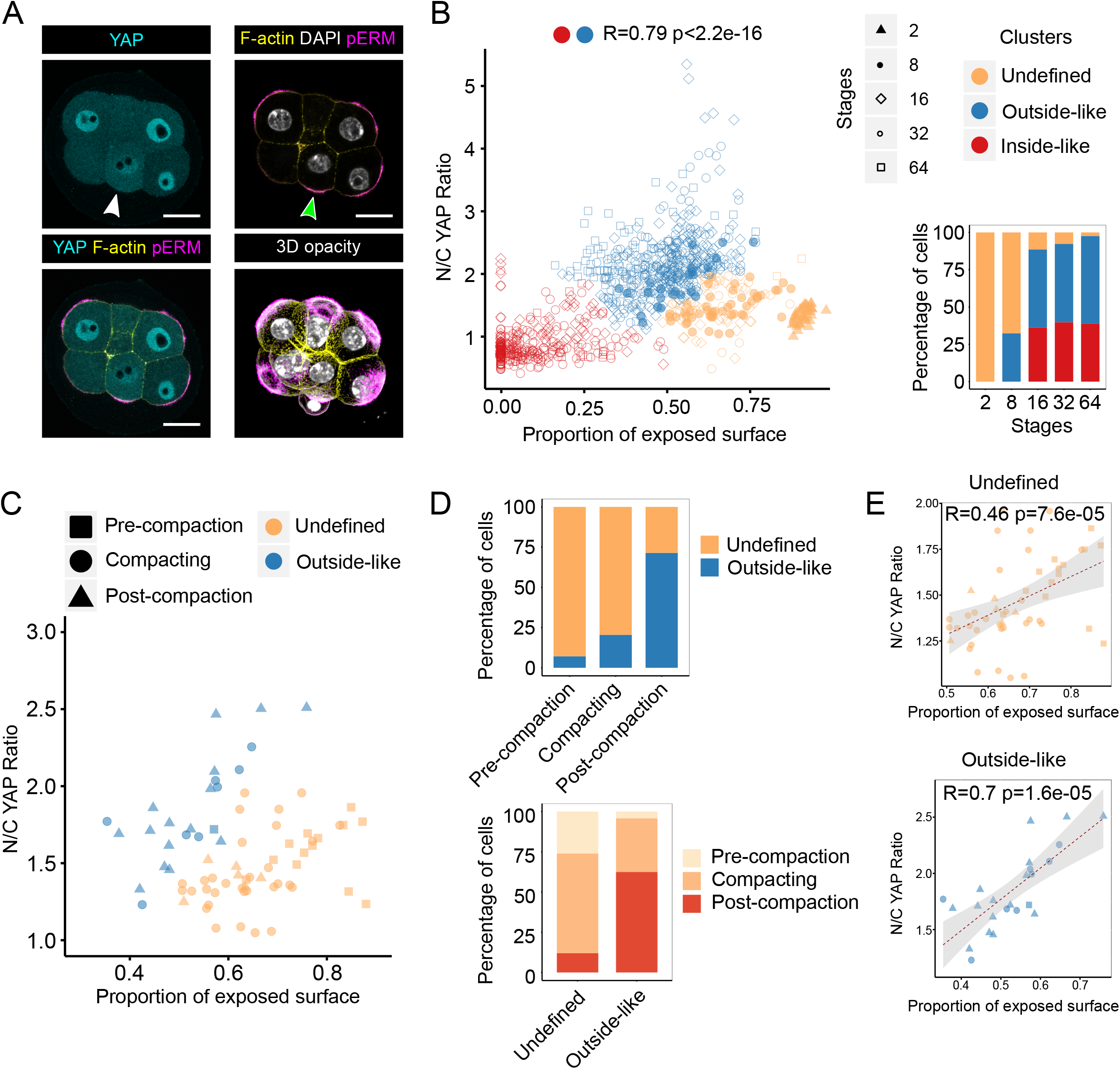
Hierarchical clustering analysis reveals the association between the proportion of exposed surface and N/C YAP ratio in compacted 8-cell embryos. A. images of an 8-cell embryo immunostained for YAP and pERM illustrating variations in N/C YAP ratio at the 8-cell stage. F-actin and nuclei were visualised using Phalloidin and DAPI respectively. White arrowhead points to a blastomere with lower N/C YAP ratio. Green arrowhead highlights the presence of apical pERM. Bottom right panel shows a 3D opacity rendering of the corresponding embryo. Scale bar: 20 μm. B. Hierarchical clustering of blastomeres across preimplantation development into three distinct clusters. Blastomeres with high N/C YAP ratio and intermediate proportion of exposed cell surface area were classified as belonging to the outside-like cluster. Blastomeres with low N/C YAP ratio and low exposed cell surface area were classified as belonging to the inside-like cluster. Finally, the remaining blastomeres, exhibiting high proportion of exposed cell surface area and intermediate N/C YAP ratio were defined as belonging to an “undefined cluster”. Dot shape indicates stage whereas colour indicates the cluster to which each blastomere belongs (Spearman, R=0.79 p<2.2e−16). Bottom right bar graph represents the distribution of blastomeres across the three clusters for each stage. C. Analysis of N/C YAP ratio and the proportion of exposed cell surface area at the 8-cell stage in not compacted, compacting and compacted embryos. D. Bar graphs representing the proportion of blastomeres from pre-compaction, compacting and post-compaction embryos in the Undefined and Outside-like clusters (top). The proportion of blastomeres from each cluster found in pre-compaction, compacting and post-compaction embryos is shown at bottom. E. Correlation between the proportion of exposed surface and N/C YAP ratio in Undefined (top) (Spearman, R=0.46 p=7.6e−05) and Outside-like 8-cell blastomeres (bottom) (Spearman, R=0.7 p=1.6e−05).

To test whether there may already be blastomeres at the 8-cell stage that display a relationship between N/C YAP ratio and the proportion of exposed cell surface area, we first tested whether blastomeres across development could be classified according to just these two characteristics. To do this, we performed unsupervised hierarchical clustering of our entire dataset using N/C YAP ratio and the proportion of exposed cell surface area as variables and found that on the basis of just these two parameters, blastomeres could be grouped into three distinct clusters (Figure 3B). One was characterised by low N/C YAP ratio and low proportion of exposed surface area, a second by high N/C YAP ratio and intermediate proportion of exposed surface area and the third by intermediate N/C YAP ratio and high proportion of exposed surface area (Figure S3A-C). Based on our observations and current knowledge, blastomeres from the first two clusters exhibited ICM- and TE-like characteristics respectively, whereas the third cluster potentially represented blastomeres in an ‘undecided’ state, as it contained large numbers of blastomeres from the 2- and 8-cell stages. We therefore named the clusters ‘inside-like’, ‘outside-like’ and ‘undefined’ respectively. Consistent with developmental trajectories, we could detect a strong positive correlation between N/C YAP ratio and the proportion of exposed surface when just the outside-like and inside-like clusters were considered together, without the undefined cluster (Figure 3B).

To determine how well the unsupervised clustering performed, we next manually categorised blastomeres from a subset of 32- and 64-cell stage embryos as inside or outside, to generate a ground truth to compare the unbiased clustering against. We found that all blastomeres manually annotated as ICM fell into the inside-like cluster (82/82) and the vast majority annotated as TE fell into the outside-like cluster (109/116, 94%). Conversely, the outside-like cluster consisted exclusively of TE cells (109/109), while the inside-like cluster consisted almost exclusively of ICM cells (82/85, 96.5%). Overall, the clustering displayed very few errors, and these lay at the interface of the different clusters (Figure S3D). The broad accuracy of the unsupervised clustering suggests that the blastomeres clustering as ‘undefined’ might represent a biologically meaningful state.

All 2-cell stage blastomeres belonged to the undefined cluster whereas only two-thirds of those from the 8-cell stage belonged to this cluster with the remaining one-third falling in the outside-like cluster (Figure 3B). Thereafter, a much lower proportion of 16- to 64-cell stage blastomeres belonged to the undefined cluster (Figure 3B and S3E). Taken together this again suggested that already at the 8-cell stage, differences were starting to emerge amongst blastomeres, that those first to become different were on a TE trajectory and that ICM cells arise only at the 16-cell stage, when cells start being found inside the embryo.

To verify whether already at the 8-cell stage blastomeres acquire the ability to sense and respond to their position, and whether this is linked to embryo maturation, we examined in more detail 8-cell embryos at different stages of compaction (Figure 3C). We categorised our 8-cell embryos based on their morphology as “Pre-compaction”, “Compacting” and “Post-compaction”. In validation of our categorisation, morphometric parameters of blastomere sphericity and proportion of exposed cell surface were significantly different in embryos post-compaction (Figure S3F and S3G).

Interestingly, the vast majority of blastomeres from post-compaction 8-cell embryos belonged to the outside-like cluster (15/21, 71.4%). In contrast, all but one blastomere from pre-compaction 8-cell embryos belonged to the undefined cluster (Figure 3D). The allocation of most blastomeres from post-compaction embryos to the outside-like cluster and from pre-compaction embryos to the inside-like cluster could not be explained by changes in YAP localisation alone, as there were no significant differences in N/C YAP ratio between blastomeres depending on the degree of embryo compaction (Figure S3H). Amongst blastomeres from the undefined cluster, there was no clear relationship between the proportion of exposed cell surface area and N/C YAP ratio (Figure 3E), suggesting that these blastomeres were not able to sense the proportion of cell membrane exposed to the outside. In contrast, 8-cell blastomeres from the outside-like cluster, which come predominantly from post-compaction embryos, exhibited a strong relationship between the proportion of exposed cell surface area and N/C YAP ratio (Figure 3E). This suggests that the position sensing machinery is in place earlier than previously thought, by the time compaction is completed.

### Embryo shape manipulation alters blastomere fate and reveals position sensing at the 8-cell stage

To test if the proportion of exposed surface area is indeed used as a measure of relative blastomere position to regulate N/C YAP ratio and ultimately, cell fate, we developed a non-invasive approach to alter the arrangement and shape of cells in the pre-compaction embryo. Our aim was to modulate the fraction of cells with high and low proportion of exposed surface area within each embryo and analyse the consequences on N/C YAP ratio. To this end, we used biocompatible hydrogels to create channels of consistent diameters in which to culture embryos. This physical constraint forced blastomere-rearrangements and caused embryos to adopt a cylindrical shape (Figure 4A and 4B and movie S2). As expected, this cylindrical distortion resulted in an increase in the overall external surface area to volume ratio of the embryo as compared to spherical controls (Figure S4A and S4B).

**Figure 4:**
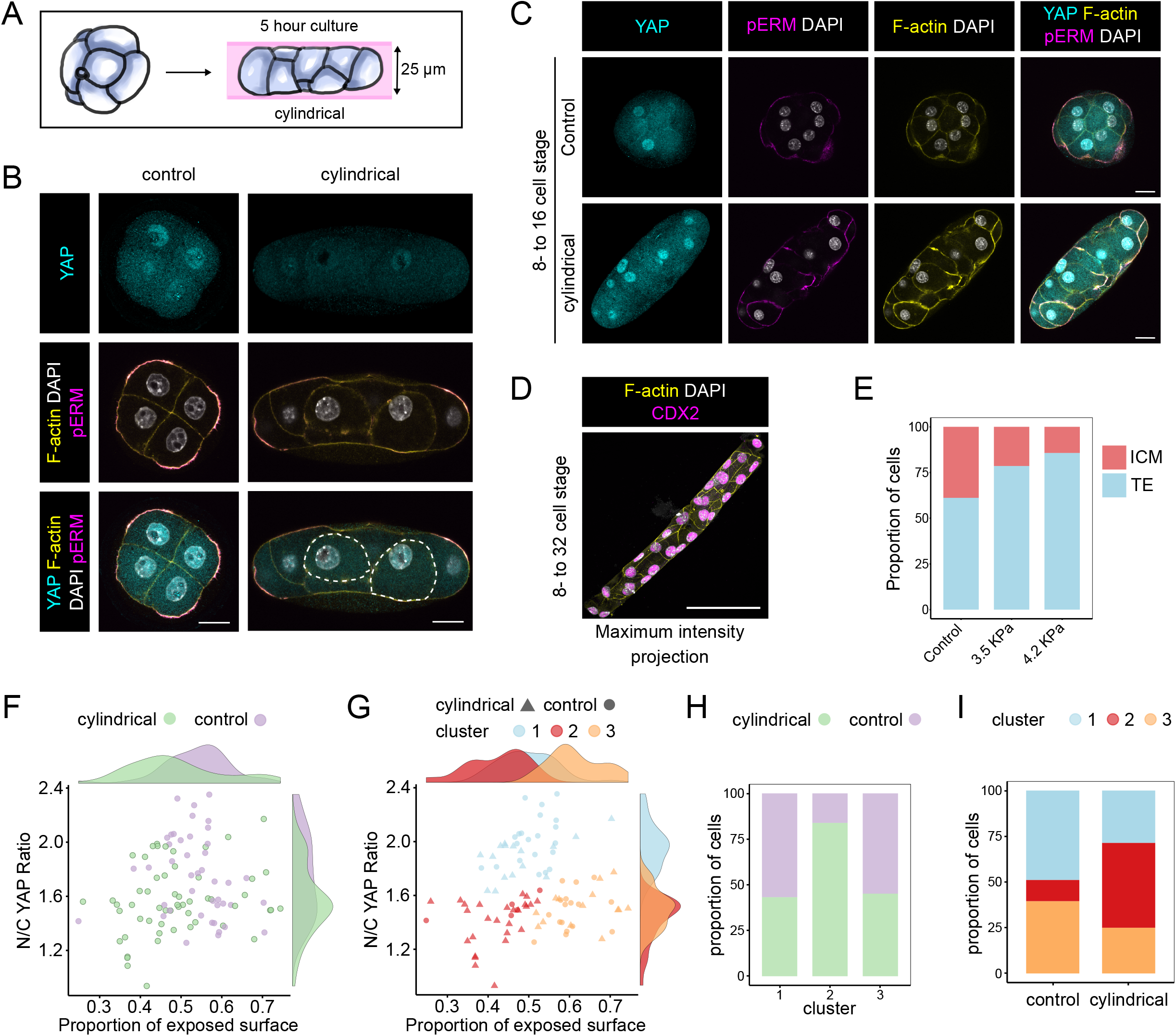
Embryo shape manipulation reveals position-sensing at the 8-cell stage. A. Diagram representing the experimental design. 8-cell embryos were inserted into 25 μm diameter channels and cultured for 5 hours. Embryos grown in channels adopted a cylindrical configuration. B. Representative images of control and cylindrical 8-cell embryos immunostained for YAP and pERM. F-actin and nuclei were visualised using Phalloidin and DAPI respectively. Scale bar: 20 μm. C. Representative images of 8-cell embryos grown to the 16-cell stage in control conditions or in 25 μm channels and subsequently immunostained using antibodies against YAP and pERM. F-actin and nuclei were visualised using Phalloidin and DAPI respectively. Scale bar: 20 μm. D. An example of an 8-cell embryo grown until the 32-cell stage in a 25 μm channel, in which all cells are forced to be on the “outside”. Note how all cells are CDX2 positive. Scale bar: 50 μm. E. Quantification of cell fates (estimated by CDX2 levels) in 16-cell embryos following two different levels of compression at the 8- cell stage, using gels of 3.5 and 4.2 KPa in stiffness respectively. Control: 121 cells; 3.5 KPa: 98 cells; 4.2 KPa: 84 cells. F. Plot showing the proportion of exposed surface and N/C YAP ratio in control (n=6 embryos) and cylindrical (n=5 embryos) embryos. Marginal density plots for control and cylindrical embryos, on the sides of the graph, show a shift in both the proportion of exposed surface and N/C YAP ratio in blastomeres from cylindrical embryos. G. Representation of the proportion of exposed surface and N/C YAP ratio in blastomeres from control and cylindrical embryos and the different clusters obtained by hierarchical clustering. H. Bar graph representing the proportion of blastomeres from control or cylindrical embryos in each cluster. I. Proportion of blastomeres from each cluster in control and cylindrical embryos.

Since our observations suggested that post-compaction, 8-cell blastomeres are on a TE trajectory (Figure 3B and 3C), we took advantage of our channel assay to address whether artificially reducing the number of cells that are inside would at later stages result in cell fate changes. When 8-cell embryos were grown inside channels until the 16-cell stage, fewer cells excluded YAP from the nucleus in comparison to the controls, suggesting that fewer ICM cells were formed (Figure 4C). To test whether this was the case, we cultured embryos from the 8- to the 32-cell-stage in channels within polyacrylamide of varying stiffness, (see methods for details) to subject them to either moderate or extreme confinement. We then stained the embryos for CDX2 to assay blastomere fate. In embryos under extreme confinement, all cells were forced to occupy the outer surface and it was possible to create embryos consisting solely of CDX2-positive cells (Figure 4D). Under milder compression, blastomeres managed to move inside, but their numbers decreased with increasing tissue deformation (Figure 4E). These experiments show that by simply preventing inside cells from arising, it is possible to derive embryos made exclusively of TE blastomeres, consistent with 8-cell blastomeres existing on a TE trajectory and with blastomere position being the main driving force behind the emergence of the ICM lineage.

Next, we used our channel assay to study position sensing at the 8-cell stage. When 8-cell embryos were inserted into channels and cultured for five hours (Figure 4B), individual blastomeres exhibited a wider range of proportion of exposed surface in comparison to blastomeres from control embryos. A few, from the tip of the cylindrical embryos, exhibited a higher than normal proportion of exposed surface while most others exhibited reduced proportion of exposed surface. This resulted in overall lower values for proportion of exposed surface in comparison to controls (Figure 4F and S4C). Consistent with the proportion of exposed surface area being used to sense the position of blastomeres, modulating the proportion of exposed surface in this way was accompanied by a significant decrease in mean N/C YAP ratio in blastomeres of cylindrical embryos (Figure 4F and S4D).

To test whether cylindrical embryos were able to generate a new population of cells simultaneously characterised by lower proportion of exposed surface and reduced N/C YAP ratio, we performed hierarchical clustering of these blastomeres on the basis of N/C YAP ratio and the proportion of exposed cell surface area. This resulted in three clusters (Figure 4G and S4E). Strikingly, Cluster 2, containing blastomeres with the lowest proportion of exposed surface area and N/C YAP ratio, was predominantly composed (83.9%) of cells from cylindrical embryos (Figure 4H) and close to half (46.4%) of the blastomeres from cylindrical embryos belonged to this cluster (Figure 4I). Together, these results show that changes to the proportion of exposed blastomere surface area causes changes in N/C YAP ratio. This tight relationship between proportion of exposed cell surface and N/C YAP ratio supports the idea that blastomeres from compacted 8-cell embryos are able to sense and rapidly respond to subtle changes in their proportion of exposed cell surface area.

### N/C YAP ratio increase from the 2- to 8-cell stage is dependent on PKC activity

What might be the molecular underpinnings whereby blastomeres sense the proportion of exposed surface? Apicobasal polarity has been shown to positively regulate nuclear localisation of YAP and offered a good candidate. Given that during compaction, all blastomeres polarise, and the exposed surface is the apical domain of the polarised blastomere, we hypothesised that blastomere might be using the extent of the apical domain when measuring the proportion of exposed surface. To test this, we used Ro-31-8220 (RO), a PKC inhibitor that leads to reduced phosphorylation of ERM proteins (pERM), a marker of the apical domain (Liu et al., 2013). Reduced apical pERM in turn leads to a decrease in apical F-actin (Liu et al., 2013), which further impairs the establishment of polarity as the actomyosin network is required for the apical localisation of the Par complex (Zhu et al., 2017). We found that embryos treated with inhibitor from the 2- to the 8-cell stage not only exhibited reduced proportion of apical to basolateral pERM (A/B pERM ratio) but were also unable to form apical actin rings (figure 5A and 5D) consistent with disruption of apicobasal polarity. Inhibition of PKC was also accompanied by a markedly reduced N/C YAP ratio (Figure 5A and 5C). Moreover, when we plotted the correlation between N/C YAP ratio and A/B pERM ratio, we could detect a moderate positive correlation (Figure 5D). Briefer 5-hour treatment of 8-cell embryos with inhibitor did not result in a comparatively strong decrease in nuclear YAP (data not shown), presumably because YAP had already had enough time to accumulate in the nucleus prior to treatment (Figure 5E). Interestingly, 5-hour RO treatment resulted in both YAP and F-actin accumulating and colocalising in small cytoplasmic aggregates (Figure 5E). High-resolution imaging showed that even in unmanipulated embryos, YAP can often be detected at cell-cell junctions (Figure 5F) suggesting that YAP can interact, most likely indirectly, with cortical F-actin, and that it remains associated with this complex but gets translocated to the cytosol upon RO treatment. Collectively, our data show that PKC activity is required from the 2- to 8-cell stage for the nuclear accumulation of YAP, suggesting that the machinery setting up apicobasal polarity could be used to measure the proportion of exposed surface area and ultimately, the position of blastomeres within the pre-implantation embryo.

**Figure 5:**
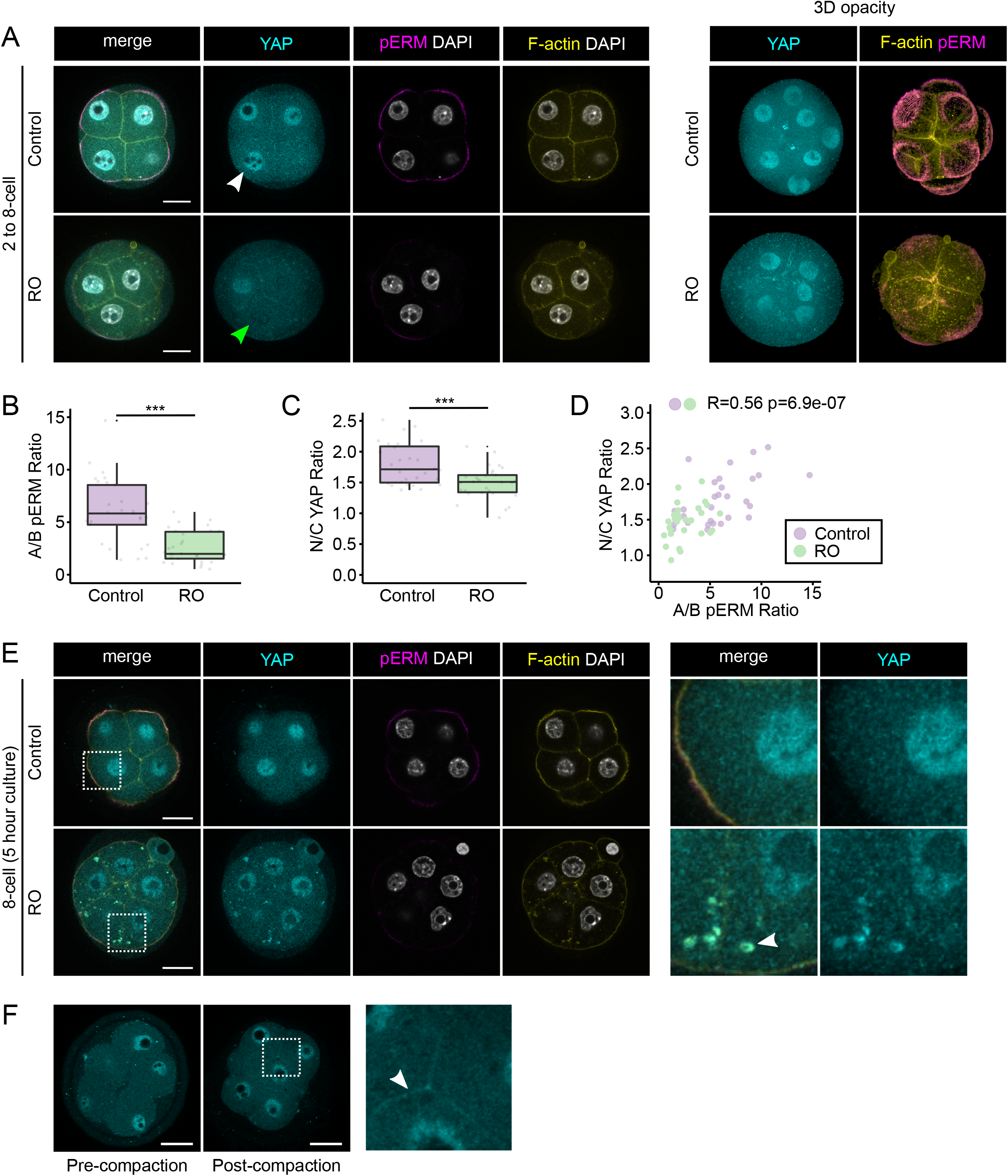
PKC regulates the nuclear accumulation of YAP from the 2- to 8-cell stage. A. Representative images of DMSO and RO-treated embryos grown *in vitro* from the 2- to the 8-cell stage immunostained for YAP and pERM. F-actin and nuclei were visualised using Phalloidin and DAPI respectively. White arrowhead points to a nucleus with high levels of YAP, whereas green arrowhead points to a nucleus with low levels of YAP. Right panel shows 3D opacity renderings of corresponding embryos. Scale bar: 20 μm. B. Boxplot showing the proportion of pERM at the apical membrane in control (n=4 embryos) and RO-treated (n=4 embryos) embryos. C. Boxplot showing the proportion of YAP in the nucleus in control and RO-treated embryos. *** p<0.001 (Kruskal Wallis test). D. Plot showing the relationship between the proportion of pERM at the apical membrane and N/C YAP ratio in control and RO-treated embryos (Spearman, R=0.56 p=6.9e−07). E. Representative images of embryos cultured for 5 hours at the 8-cell stage in the presence of either DMSO or RO and subsequently immunostained for YAP and pERM. F-actin and nuclei were visualised using Phalloidin and DAPI respectively. Right panel shows magnification of the areas surrounded by dashed outlines. The white arrowhead points to cytoplasmic puncta of F-actin and YAP. Scale bar: 20 μm. F. Representative images of pre- and post-compaction 8-cell embryos immunostained for YAP and showing blastomeres with comparable N/C YAP ratios. The panel at right shows a high-magnification image of the boxed area. The white arrowhead points at YAP localised at cell-cell junctions.

## DISCUSSION

We have discovered that, following a PKC-dependent accumulation of YAP in the nucleus during compaction at the 8-cell stage, blastomeres start exhibiting a close relationship between their proportion of exposed surface and the proportion of YAP in their nucleus and cytoplasm. This suggests that blastomeres, following polarisation, can sense their relative position within the embryos and transfer this information to the nucleus by modulating the subcellular localisation of YAP. To demonstrate the position-sensing ability of blastomeres within the embryo, we used a novel approach that, as opposed to the more traditional use of chimeras or dissociated blastomeres, does not alter the number of cells or the structural integrity of embryos (Leonavicius et al., 2018). By inserting embryos within cylindrical channels of defined diameter, we were able to alter embryo shape and the relative position of blastomeres. Using this method, we also showed that blastomeres of the 8-cell embryo are on a TE trajectory because, when blastomere internalisation was prevented, TE fate was favoured. This demonstrates the crucial role of blastomere internalisation in the emergence of the ICM lineage.

What mechanism might blastomeres use to measure the proportion of their surface that is exposed to the exterior? Polarisation at the 8-cell stage is the first event generating asymmetries within blastomeres from a structural point of view (Ducibella et al., 1977; Louvet et al., 1996; Vinot et al., 2005; Ziomek and Johnson, 1980). Recent work has advanced our understanding of the signalling pathways involved in the polarisation process (Zhu et al., 2017). The actomyosin contractility machinery is recruited to cell contact-free membrane in a chain of events initiated by phospholipase C (PLC). Ultimately, this leads to the recruitment of Par6 and aPKC and the establishment of apicobasal polarity. Incidentally, aPKC activity has been shown to be required for the phosphorylation of ERM proteins, which is thought to be important during compaction (Dard et al., 2004; Liu et al., 2013). Importantly, the organisation of the contractility machinery at the cell cortex enables the generation of the forces driving compaction (Maître et al., 2015). This body of work highlights one of the major outcomes of compaction: the creation of two different membrane domains, the apical and basolateral domains, with different organisation, protein complexes and contractile properties. Several of our observations suggest that YAP may be able to localise to these different membrane domains. First, we can detect YAP at cell-cell junctions, along the basolateral domain, in 8- cell embryos. Second, when PKC was inhibited in 8-cell embryos, YAP accumulated in cytoplasmic puncta where it colocalised with F-actin. Since PKC inhibition reduces the phosphorylation of apical ERM and disrupts apical F-actin organisation, this suggests that YAP can interact, presumably indirectly, with apical actin. Together, these observations suggest a mechanism whereby YAP localised at the apical and basolateral domains are regulated differently. The balance between inhibitory signal at the basolateral membrane and activating signal at the apical membrane might provide the cell with a measure of the proportion of exposed cell surface area. Currently, amongst YAP negative regulators, only AMOT has been shown to distribute asymmetrically in the polarised cells of the preimplantation embryo (Hirate et al., 2013; Leung and Zernicka-Goetz, 2013). In other epithelial tissues, various reports suggest that YAP may be differentially regulated by complexes that localise to different membrane domains. For example, adherens junctions have been proposed to be able to sequester YAP through α-Catenin (Schlegelmilch et al., 2011), whereas YAP can associate with and be positively regulated by the ASPP2/PP1 module at tight junctions (Royer et al., 2014). Different protein complexes within the same membrane domain can also have different consequences on YAP activity. For instance, disruption of adherens junction or basolateral complexes differentially regulates YAP activity (Yang et al., 2015). Hippo interactome studies reveal that YAP interacts, directly or indirectly, with various components of not only the apicobasal polarity complexes but also planar polarity molecules and other proteins associated with the plasma membrane (Couzens et al., 2013; Hauri et al., 2013; Wang et al., 2014). It will therefore be important to determine which of those play a role in the preimplantation embryo to regulate the subcellular localisation of YAP in the context of position sensing.

We have found that a position sensing mechanism emerges during compaction. Following compaction, during the next round of cell divisions, the first inner cells will be formed. However, to this day, it remains unclear how precisely this is achieved (White et al., 2018). It has been suggested that cell fate decisions operate differently from the 8- to 16-cell stage (Anani et al., 2014) and from the 16- to 32-cell stage respectively (Hirate et al., 2013), with asymmetric cell divisions making a more important contribution during the 8- to 16-cell transition. Asymmetric cell divisions are thought to determine fate via the asymmetric inheritance of the apical domain (Korotkevich et al., 2017). However, clear asymmetric divisions rarely occur (Samarage et al., 2015) and in many cases, the apical domain seems to disassemble when blastomeres divide before being re-established de novo after cytokinesis (Zenker et al., 2018). Our results indicate that blastomere position within the embryo, as a result of *either* the mode of division or movement of cells, is ultimately what defines cell fate, consistent with cell division angle not determining fate (Watanabe et al., 2014). Instead, we propose that changes of global embryo geometry via cell internalisation drive the formation of the ICM, which is consistent with the important role of apical constrictions in cell internalisation to form the ICM (Anani et al., 2014; Samarage et al., 2015). Since we see a relationship between the relative amounts of YAP in the nucleus and the proportion of exposed surface, it raises the question whether nuclear YAP levels themselves can influence cell internalisation. It seems plausible that after cell division at the 8-cell stage, due to increased cell crowding, some cells exhibit a lower proportion of exposed surface by chance, leading to lower relative nuclear YAP levels. It is tempting to speculate that this triggers a positive feedback loop between reduced proportion of exposed surface and the progressive nuclear exclusion of YAP, leading to cell internalisation. This hypothesis is supported by work in *Drosophila*, showing the importance of the Hippo pathway in regulating apical domain size and apical complexes (Genevet et al., 2009; Hamaratoglu et al., 2009). Ultimately, identifying the genes that are regulated by the Hippo pathway during preimplantation development will help shed light on these challenging questions. Together, our results highlight the truly regulatory nature of the mouse preimplantation embryo to adapt and integrate global geometry changes into cell fate decisions.

## MATERIALS AND METHODS

### Mouse husbandry and embryo collection

Mice were housed in a 12-hour dark, 12-hour light cycle. CD1 females (Charles River, England) were crossed with C57BL/6J males (in house) to obtain stage specific embryos. Noon of the day of finding the mating plug was defined as 0.5 day post coitum (dpc). Most experiments were performed using flushed embryos from natural mating. For super-ovulations, 8-week old CD1 females were given intra-peritoneal injections of 5 IU of pregnant mare serum gonadotropin (PMSG) followed by 5 IU of human chorionic gonadotropin (hCG) 48 h later and were mated with C57BL/6J males. Embryos were flushed using M2 medium at the indicated stages (Sigma M7167). All experimental procedures complied with Home Office regulations (Project licence 30/3420) and were compliant with the UK animals (Scientific Procedures) Act 1986 and approved by the local Biological Services Ethical Review Process.

### Human embryo collection

Human embryos were donated from patients attending the Oxford Fertility with approval from the Human Fertilization and Embryology Authority (centre 0035, project RO198) and the Oxfordshire Research Ethics Committee (Reference number 14/SC/0011). Informed consent was attained from all patients. Embryos were fixed in 4% paraformaldehyde, washed twice and kept in 2 PBS containing 2% bovine serum albumin (PBS-BSA) at 4°C until they were used for immunostaining.

### Wholemount immunostaining

Following fixation with 4% PFA for 15 minutes, embryos were washed twice in 2% PBS-BSA. Embryos were then permeabilized with PBS containing 0.25% Triton X-100 (PBS-T) for 15 minutes and subjected to two washes in 2% PBS-BSA at room temperature. Embryos were then placed in blocking solution for 1 hour (3% BSA, 2.5% donkey serum in PBS containing 0.1% Tween). Incubation with primary antibodies took place overnight at 4°C in a humidified chamber. The following primary antibodies were used in this study and diluted in blocking solution at the indicated concentrations: mouse-anti-YAP, 1/100 (Santa Cruz Biotechnology, sc-101199), rat-anti-E-cadherin, 1/100 (Sigma, U3254), rabbit anti-pERM, 1/200 (Cell Signaling, 3141), rabbit anti-CDX2, 1/100 (Cell Signaling, 3977). The next day embryos were washed three times in 2% PBS-BSA for 15-20 minutes. Embryos were then incubated with secondary antibodies and Phalloidin (1/100 in blocking solution for 1 hour). The following reagents were used: Alexa fluor 555 donkey-anti-mouse (Invitrogen, A-31570), Alexa fluor 647 goat-anti-rat (Invitrogen, A-21247, Alexa fluor 488 donkey-anti-rabbit (Invitrogen, A21206), Phalloidin-Atto 488 (Sigma, 49409), Phalloidin–Atto 647N (Sigma, 65906). After another three washes of 15-20 minutes in 2% PBS-BSA, the embryos were mounted in 8-well chambers in droplets consisting of 0.5μl Vectashield with DAPI (Vector Laboratories) and 0.5 μl 2% PBS-BSA. Embryos were transferred between solutions by mouth-pipetting. All incubations took place at room temperature, unless stated otherwise. After mounting the embryos were kept in the dark at 4°C until they were imaged.

### Confocal Microscopy

Embryos were imaged on a Zeiss LSM 880 confocal microscope, using a C-Apochromat 40x/1.2 W Korr M27 water immersion objective. Laser excitation wavelengths were 405, 488, 561 and 633 nanometres depending on specific fluorophore. Embryos were imaged using a 1.5x zoom at a resolution of 512×512 pixels and 8-bit depth Z-stacks of entire embryos were acquired at a 1 μm interval using non-saturating scan parameters.

### Embryo culture

Embryos were cultured in organ culture dishes in 500 μl pre-equilibrated Evolve medium (Zenith Biotech) at 37°C and 5% CO2 for the indicated amount of time. The PKC inhibitor Ro-31-8220 was diluted in DMSO and used at 2.5 μM (1/2000 dilution). The same amount of DMSO was used in control cultures.

For cylindrical embryo cultures, channels were formed by casting a 5% (which corresponded to approximately 4.2kPa stiffness) acrylamide hydrogel (containing 39:1 bisacrylamide) around 25 μm wires within the confinement of a two-part mould (10×10×1mm). In milder compression experiments, the amount of acrylamide/bisacrylamide was reduced to create softer gels of approximately 3.5kPa stiffness (Tse and Engler, 2010). Ammonium persulphate (0.1%) and TEMED (1%) were added to polymerize polyacrylamide. The wires were then removed to form cylindrical cavities within hydrogel pieces, which were cut to roughly 3×3×1mm blocks for easier manipulation during embryo insertion. The hydrogels were carefully washed and equilibrated in embryo culture media at 37°C and 5% CO2 over-night. The embryos were then inserted into the channels using a glass capillary with a diameter slightly larger than the embryo itself. It was used to stretch the hydrogel channel before injecting the embryos and letting the channels relax and deform the embryos. Cell viability in channels had previously been assessed without any noticeable difference with control embryos (Leonavicius et al., 2018). At the end of the experiments, embryos were fixed inside the hydrogels with 4% PFA for 20 minutes. Once fixed, embryos were then removed from the hydrogel channels and immunostained in parallel to the controls.

### Segmentation and image analysis

Manual segmentation of confocal data was done using Imaris v6.3 (Bitplane). Cell and nucleus outlines were drawn using a Wacom Cintique 21UX tablet display to create a 3D surface of each blastomere membrane and nucleus using the contour surface function. Information about geometry (sphericity, surface area, volume, oblate and prolate) and signal intensity within each compartment could then be exported. Information about blastomere exposed, contact and junctional surface areas was obtained by considering surface proximities and was automated using an in-house developed software (Javali et al., 2017; Leonavicius et al., 2017). Signal intensity around these defined membrane domains could then be extracted. Dividing cells were excluded from the analysis as their geometry parameters were widely different to non-dividing cells and their nuclear envelope disassembled. Imaris files of segmented embryos will be made available on request.

### Clustering and Statistical analysis

Figures and diagrams were assembled and created using the free and open source software, Inkscape and Krita. All statistics and graphs were done using RStudio and R. Graphs were produced using several packages, including ggplot2 and ggpubr. For statistical analysis, normality of the data was first assessed using visualisation tools and statistical tests (Shapiro-Wilk normality test). When the data was normally distributed, we used Analysis of variance (ANOVA), followed by post hoc comparisons using the Tukey HSD test when comparing more than two conditions. Otherwise, the Kruskal-Wallis test was used, followed by post hoc comparison using the Dunn test. To test the correlation between two variables, the Spearman method was used when the two variables were not normally distributed. The Corrplot package was used to create a correlation matrix of the different variables in the preimplantation dataset. To define the N/C YAP Ratio threshold between cells with high and low YAP ratio by k-means clustering, the data was standardized, and distance measures were obtained using the Euclidean method. For hierarchical clustering analysis, the dynamicTreeCut function was used to determine the ideal number of clusters. The variables were scaled, and the distance matrix was produced using the Euclidean method. The Ward method was used to perform the clustering.

## Acknowledgements

We thank Dr Jonathan Godwin for his help with the superovulation procedures. This work was funded through Wellcome Senior Investigator Award 105031/C/14/Z to SS.

## Author Contributions

C.R., K.L. and S.S. conceived and designed the experiments. C.R., K.L., A.K., D.F., K.N. and C.G. conducted the experiments. C.R., K.L., A.K. and S.S. analysed the data. C.R. performed the statistical analyses. A.V., C.J., T.C., K.C. and C.G. organised the collection of human embryos. C.R. and S.S. wrote the manuscript.

---

**Figure S1:**
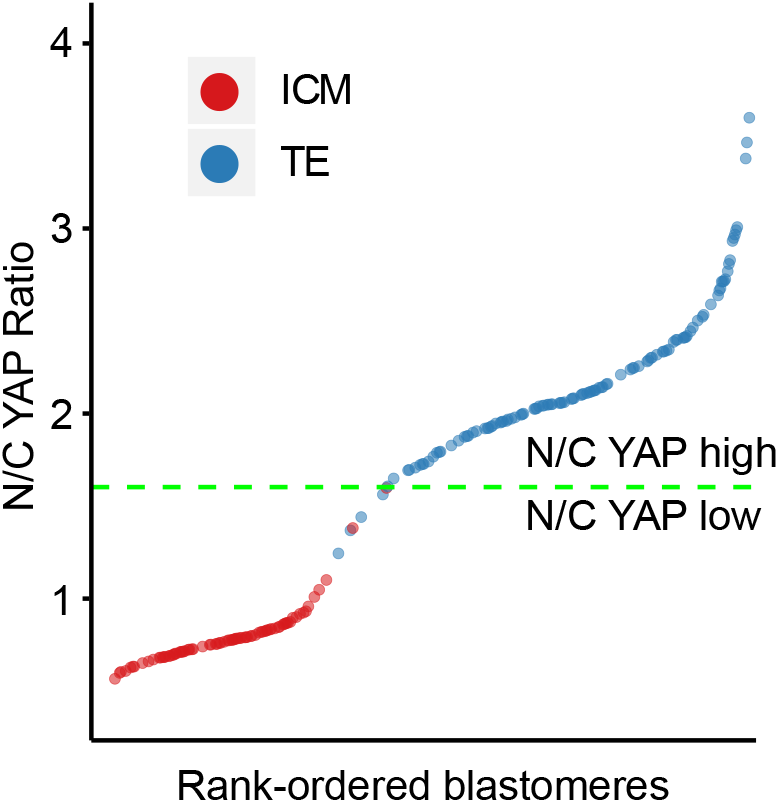
Unbiased separation of blastomeres with low and high N/C YAP ratio using K-means clustering. Blastomeres from 32- and 64-cell embryos were ranked by ascending N/C YAP ratio. The green dashed line represents the threshold value separating high and low N/C YAP ratios, as defined by the K-means algorithm. To validate the unsupervised clustering, ICM (red) or TE (blue) identity was manually assigned to blastomeres based on their position.

**Figure S2:**
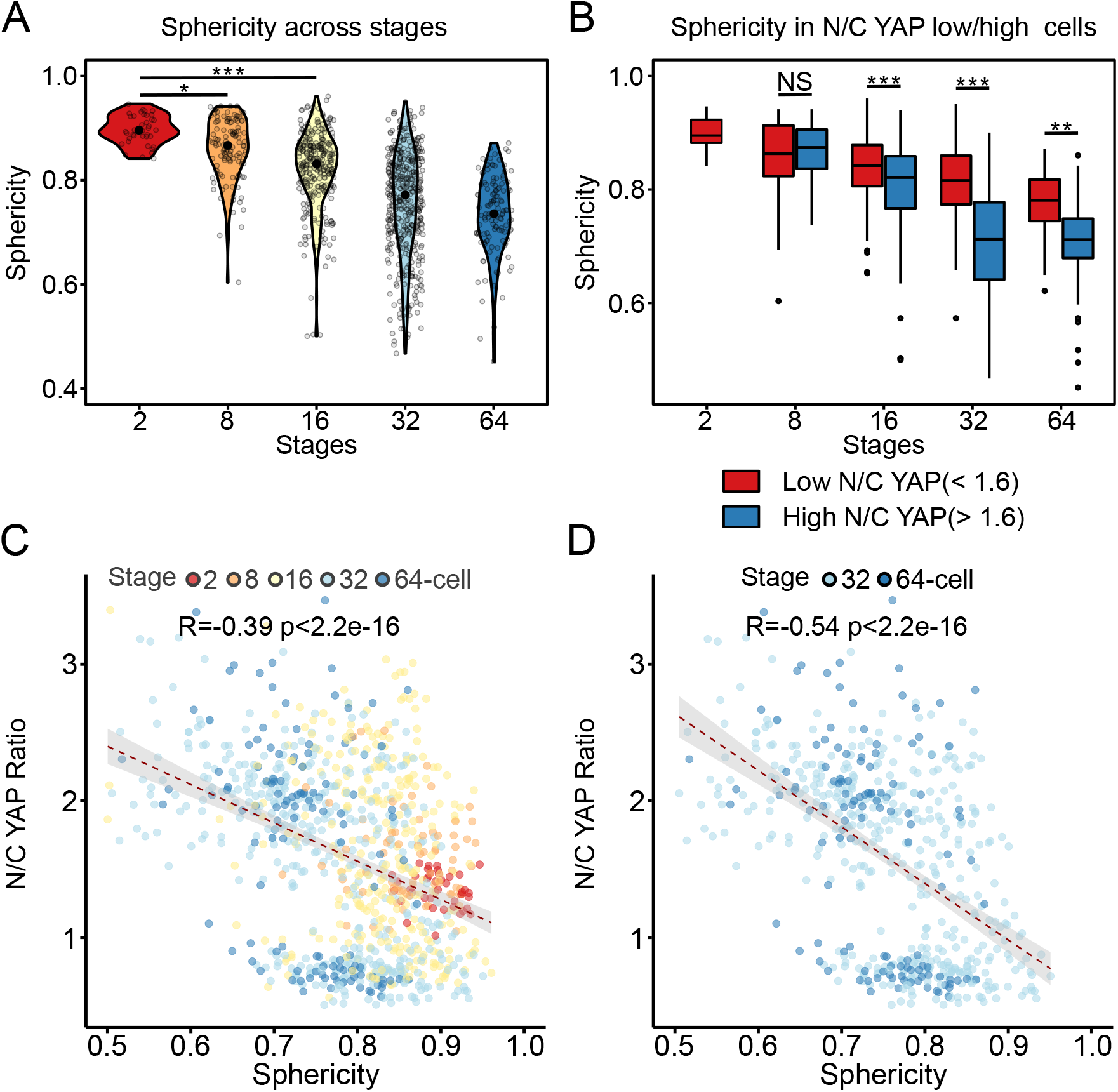
Sphericity is a poor predictor of N/C YAP ratio. A. Sphericity across developmental stages. Mean Sphericity for each developmental stage is represented as a black dot. B. Sphericity of blastomeres with either low or high N/C YAP ratio across developmental stages. NS: not significant. NS: not significant. * p<0.05, ** p<0.01, *** p<0.001 (Kruskal Wallis test followed by Dunn’s test). C. Correlation analysis between sphericity (shape) and N/C YAP ratio throughout preimplantation development (Spearman, R=−0.39 p<2.2e-16). D. Correlation analysis between sphericity and N/C YAP ratio from the 32-cell stage during preimplantation development (Spearman, R=−0.54 p<2.2e-16).

**Figure S3:**
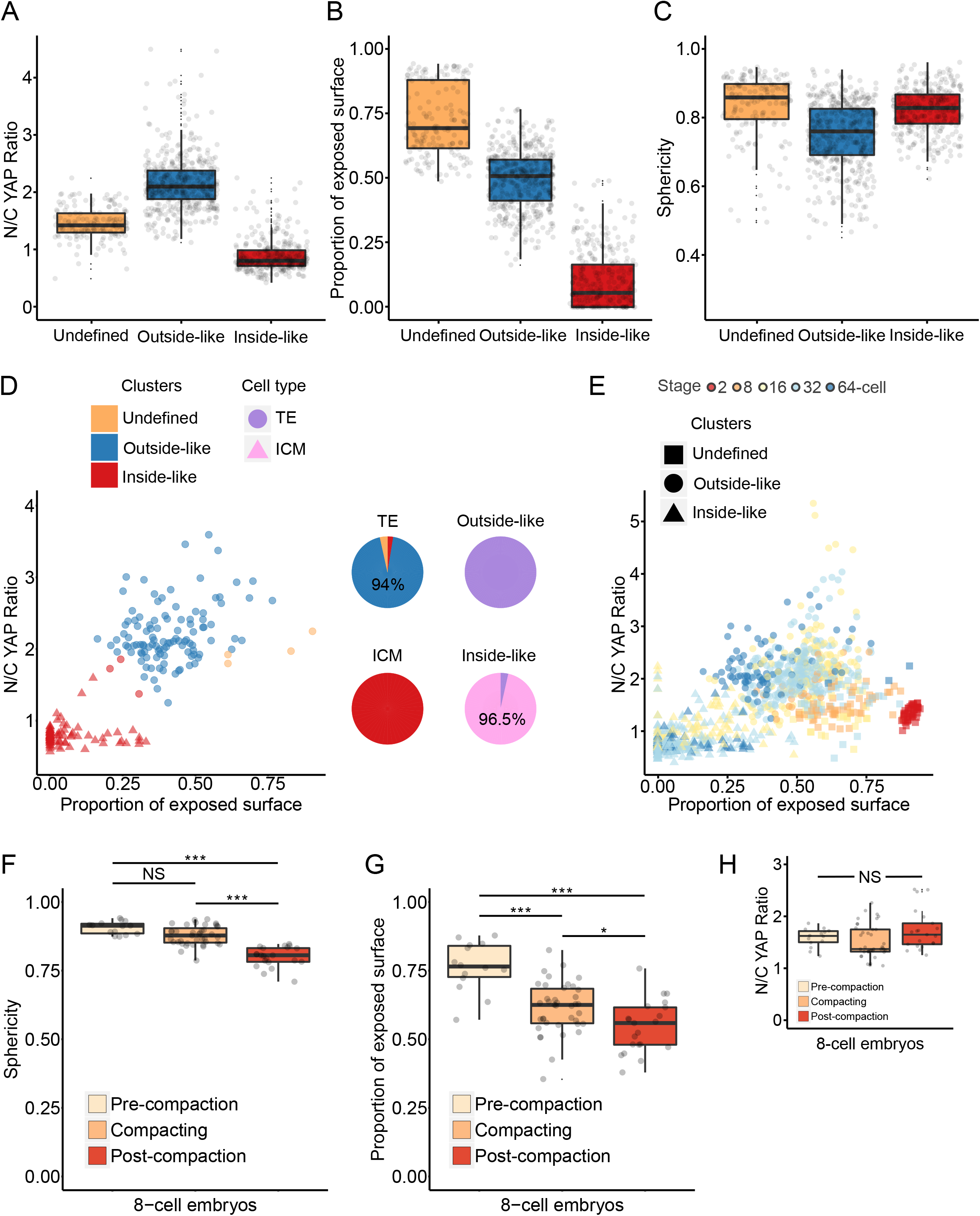
Validation of clusters obtained by hierarchical clustering. A. N/C YAP ratio of individual blastomeres across development in undefined, outside-like and inside-like clusters B. Proportion of exposed surface of individual blastomeres in undefined, outside-like and inside-like clusters. C. Sphericity of individual blastomeres in undefined, outside-like and inside-like clusters. D. Unsupervised hierarchical clustering of blastomeres from 32- and 64-cell embryo (see results for description of the three cluster types). ICM (pink) or TE (light purple) identity was manually assigned to blastomeres based on their position to validate the clustering. The pie charts display the proportion of different types of blastomeres that make up each category. All Outside-like and the majority of Inside-like blastomeres were TE and ICM respectively. E. Scatter plot of blastomere proportion of exposed surface and N/C YAP ratio in blastomeres across development (dot colour represents stages) with information on the clusters they belong to (dot shape).F, G and H. Sphericity, proportion of exposed surface area and N/C YAP ratio of individual blastomeres in pre-compaction (n=2 embryos), compacting (n=5 embryos) and post-compaction (n=3 embryos) embryos. NS: not significant. *** p<0.001, * p<0.05 (Kruskal Wallis test followed by Dunn’s test).

**Figure S4:**
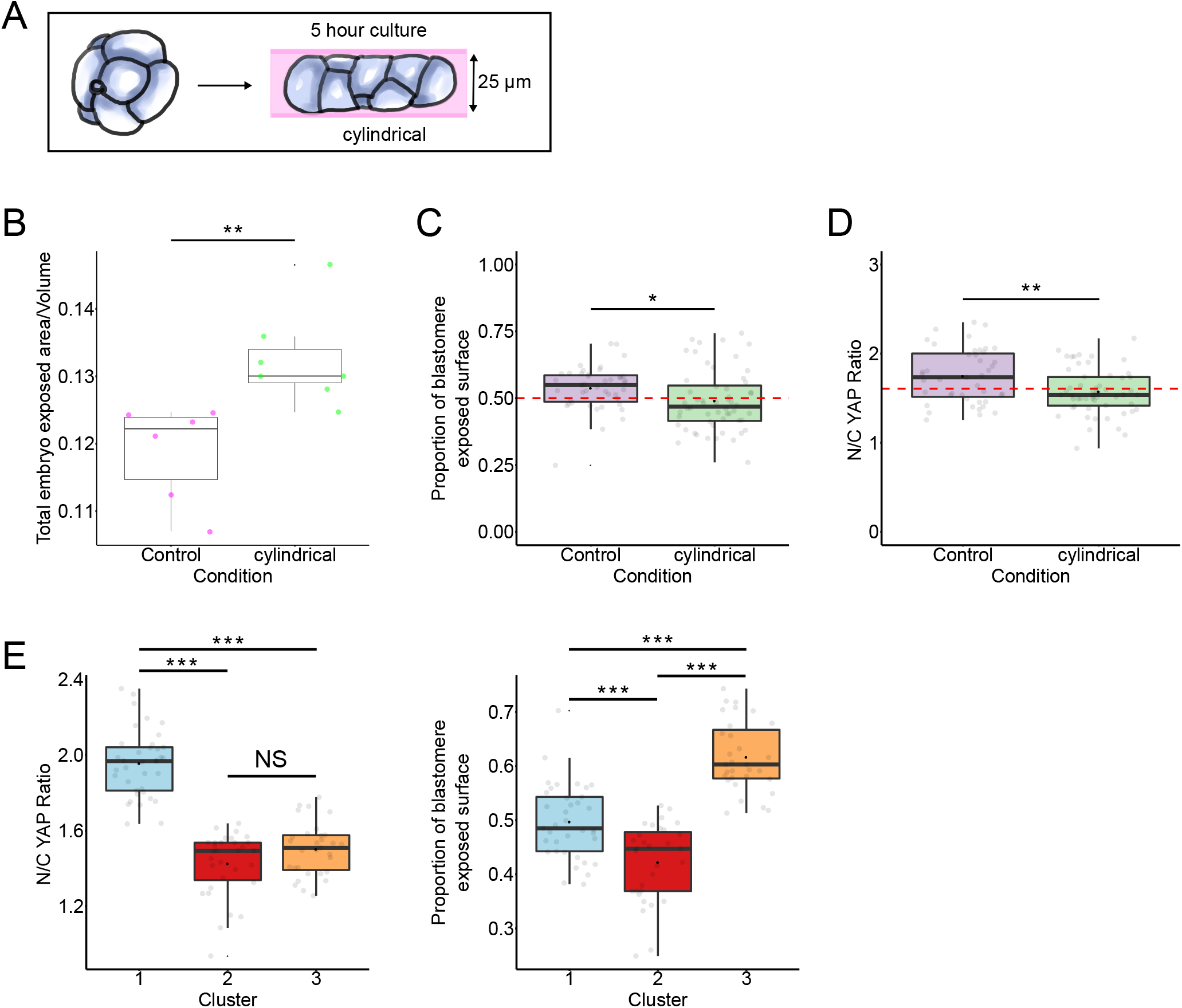
Effect of embryo shape manipulation at the 8-cell stage on embryo and blastomere geometry. A. Diagram representing the experimental design. 8-cell embryos were inserted into 25 μm diameter channels and cultured for 5 hours. Embryos grown in channels adopted a cylindrical configuration. B. Boxplot representing the surface area to volume ratio of control (n=6 embryos) and cylindrical (n=5 embryos) embryos. ** p<0.01 (T-test). C. Boxplot of the proportion of exposed surface in control and cylindrical embryos. D. Boxplot of N/C YAP ratio in control and cylindrical embryos. E. N/C YAP ratio and proportion of exposed surface in the different clusters obtained from control and cylindrical embryos based on N/C YAP ratio and the proportion of exposed surface area. NS: not significant. * p<0.05, ** p<0.01, *** p<0.001 (ANOVA followed by the Tukey HSD test)

